# Bioinformatic analysis deciphers the molecular toolbox in the endophytic/pathogenic behavior in *F. oxysporum* f. sp. *vanillae* - *V. planifolia* Jacks interaction

**DOI:** 10.1101/2021.03.23.436347

**Authors:** MT Solano De la Cruz, EE Escobar Hernández, JA Arciniega González, RP Rueda Zozaya, J Adame García, M Luna – Rodríguez

**Author notes:** These authors contributed equally to this work (MT Solano De la Cruz, EE Escobar Hernández).

## Abstract

**Background:** *F. oxysporum* as a species complex (FOSC) possesses the capacity to specialize into host-specific pathogens known as *formae speciales*. This with the help of horizontal gene transfer (HGT) between pathogenic and endophytic individuals of FOSC. From these pathogenic *forma specialis, F. oxysporum* f. sp. *vanillae* (*Fov*) is the causal agent of fusarium wilt producing root and stem rot (RSR) and is positioned as the main phytosanitary problem in vanilla plantations worldwide. Nonetheless, the origin of this forma speciales and the behavioral genetics dictating the endophytic/pathogenic *Fusarium* lifestyles still unknown. To elucidate the underlying molecular mechanisms that establish these behaviors we analyzed the RNA-seq libraries of two-times frames of vanilla-Fov interactions.

**Results:** Our analyses identified the sets of transcripts corresponding to *Fov* pathogenic strain JAGH3 during the two-times frames of the infection as the sets of the transcripts belonging to endophytic *Fox* in vanilla. Functional predictions of *de novo* annotated transcripts as the enriched GO terms with the overrepresented metabolic pathways, allowed us to identify the processes that establish the pathogenic lifestyle in *Fov* being virulence, hypervirulence, sporulation, conidiation, necrosis and fusaric acid related genes with the carbohydrates, amino acids, proteins, glycerophospholipids and autophagy metabolic pathways that are key regulators of spores germination and pathogenicity establishment as the underlying mechanisms behind this behavior. As the absence of these were found in the vanilla endophytic *Fox*.

**Conclusions:** This work reveals the main players of the behavioral genetics in pathogenic *Fov*/endophytic *Fox* in *V. planifolia* Jacks. Its pathogenic strategy allows *Fov* to infect in a SIX genes-independent manner. As the other pathogenic elements found in this study could be explained by the presence of pathogenicity islands and genomic regions associated with supernumerary chromosomes in *Fov*. These play a central role as carriers of genes involved with pathogenic activity and could be obtained through HGT.

## Background

The main phytosanitary problem affecting vanilla production worldwide is RSR caused by the phytopathogenic fungus *Fov* (Pinaria et al., 2015; Tombe et al., 1994, Thomas et al., 2002). This fungus has been reported in vanilla growing regions around the world; however, its origin and evolution still poorly understood (Pinaria et al., 2015). Pinaria et al., (2010) reported that the management of the diseases caused by *Fox* (*F. oxysporum*) eventually leads to obtaining resistance (Fravel et al., 2003), however in the vanilla crop, the plant material continues to be highly susceptible. The isolates, genetically and morphologically different from *Fox*, have been classified as special forms depending on the host from which they were obtained, these special forms establish FOSC (Booth, 1971; Michielse and Rep, 2009). Within FOSC exist a broad genomic plasticity, diversifying into different special forms with distinct genomic composition, also having the presence of supernumerary chromosomes among the special forms of FOSC, in fact it has been reported that the presence of certain supernumerary chromosomes, like chromosome 14, bestow pathogenicity to non-pathogenic strains (Ma et al., 2010). Previously it was stated that within each special form, the same genomic structure and the same evolutionary origin were shared (Taylor et al., 1999b) allowing an adequate taxonomic classification based on the sequence of some specific genes (Pinaria et al., 2015; O’Donnell et al., 1998; Taylor et al., 1999a); however, the F. oxysporum f sp. cubense and F. oxysporum f. sp. vanillae have multiple independent origins (O’Donnell et al., 1998; Flores-de la Rosa et al. 2018) showing the genomic plasticity within FOSC. In fact, Pinaria et al., (2015), mentioned that most of the special forms do not have a monophyletic origin, which suggests significant differences in their genomic structure. Also, the genetic diversity of *Fov* has been demonstrated by presenting multiple vegetative compatibility groups (Tombe et al., 1994; Pinaria, 2010), as well as several different fingerprint haplotypes (Pinaria et al., 2015). Based on these vegetative compatibility groups (Elias et al., 1993), it was proposed that the evolutionary origin of *Fov* is complex and not monophyletic having Mexico as the possible center of origin. *Fov* has been isolated as an endophyte from vanilla stems without any external or internal symptoms and it also shown to be pathogenic in greenhouse trials (Liew et al., 2008).

In 2019, our research group reported the increased expression of gene transcripts associated with ribosomal proteins during the early stages of infection caused by *Fov* in *V. planifolia* Jacks root (Solano De la Cruz et al., 2019). In this work, we found that the RNA libraries in the transcriptome under study contain readings corresponding to *Fov*, these were aligned against the reference genome of *Fox*; Fol 4287 (*Fusarium oxysporum* f. sp. *lycopersici*). Given those corresponding to *Fov*, in each of the libraries analyzed being in a range of around 5 to 39%, with the content of the lowest readings in the controls with *Fox* readings compared to the treatments. Here, we detailed the results of the alignment of *Fov* JAGH3 strain readings to the reference genome Fol 4287, its annotation and the identification of *Fov* transcripts, present during the *Fov* infection process in vanilla, at two-time frames 2 days post-inoculation (dpi) and 10 dpi. This being the first report of coding sequences corresponding to the proteins with vital functions in the fusarium wilt caused by *Fov* in vanilla on a large scale. The foregoing as an effort in determining the transcripts present in the JAGH3 strain, which are not present in *Fox* lineages with endophytic lifestyle in the vanilla root, which could explain its pathogenicity indirectly, as a first approach to obtaining the *Fov* transcriptomic and genomic data.

## Results

The quality analysis of the libraries corresponding to the *V. planifolia* Jacks transcriptome in response to the infection caused by *Fov*, reported by our working group in 2019 (Solano De la Cruz et al., 2019), was again accomplished. Later, the alignment of the libraries against the Fol 4287 genome was performed, obtaining only the readings that correctly aligned against it. About the control libraries, in each of the replicas at 2 dpi 7158305, 7249959, and 5741789 aligned readings were acquired. While at 10 dpi 6458564, 6730883, and 6528340 readings were obtained. In 2 dpi, a total of 10917618 aligned readings were collected and at 10 dpi 9397911, 10190151, and 10190151, aligned readings were retrieved. Next, we filtered the alignments results by alignment quality and sorted them by genomic coordinates.

To be able to identify the genes of the 2 dpi and 10 dpi controls and treatments, we carried out intersections of the filtered readings with the annotation of Fol 4287 genome utilizing the genomic coordinates. Table 1 presents the number of genes identified corresponding to each one of them. Finally, we determined the annotated genes that are shared between treatments and the treatment-specific ones. From the annotated transcripts we used their identifiers to retrieve their protein sequences from Biomart-Ensembl Fungi database for each of the libraries. For each of the obtained coding protein data sets. After that, a *de novo* functional annotation was performed using InterproScan 5 software (Jones et al., 2014).

**Table 1.** Summary of the filter analysis of the reads aligned with the reference genome of Fusarium oxysporum f. sp. vanillae Fol 4287, contrasting the transcripts present in the treatment libraries with the JAGH3 strain vs the controls and at 2 dpi and 10 dpi.

| Dataset to filter | Number of genes | Dataset used as filter | Number of genes | Generated dataset | Number of genes obtained | Number of genes filtered |
| --- | --- | --- | --- | --- | --- | --- |
| 2 dpi treatment | 6174 | 2 dpi control | 5455 | Genes involved in infection at 2 days | 3045 | 3129 |
| 10 dpi treatment | 13092 | 10 dpi control | 5765 | Genes involved in infection at 10 days | 7734 | 5358 |
| Genes involved in infection at 2 days | 3045 | Genes involved in infection at 10 days | 7734 | Unique genes of early infection | 1334 | 1711 |
| Genes involved in infection at 10 days | 7734 | Genes involved in infection at 2 days | 3045 | Unique genes of late infection | 6023 | 1711 |

Here, we were able to determine a collection of specific *Fov* transcripts, which were found only in the vanilla roots that were inoculated with the JAGH3 strain being directly related to the pathogenicity of *Fov* in *V. planifolia* Jacks (see in supplementary material the files 10_dpi_treatment_genes_list.txt and 2_dpi_treatment_genes_list.txt).

### Aligned genomic regions revealed the participation of supernumerary chromosomes in the establishment of the pathogenicity of *Fov* in *V. planifolia* Jacks

In figure 1 we observed the *Fox* transcripts corresponding to the control treatment, and their location on the different chromosomes in the Fol 4287 reference genome. Interestingly, we observed that many of the transcripts corresponding to endophytic *Fox* were mainly located in the chromosomes that establish the core of the reference Fol 4287 genome and not in the supernumerary chromosomes where we observed the presence of few transcripts. In the control treatment at 2 dpi, we observed the presence of transcripts corresponding mainly to chromosomes 1, 2, 4, 5, 7, 8, 9 and 10 with the presence of some transcripts in chromosomes 3, 6, 11, 12, 13 and 14; while they did not present any transcript related to chromosome 15. In the control treatment at 10 dpi, the most represented chromosomes are 1, 2, 4, 5, 7, 8, 9 and 10, while some transcripts related to chromosome 3, 11, 12 and 13 were found; Interestingly, transcripts related to chromosomes 6, 14, and 15 were absent.

**Figure 1.**
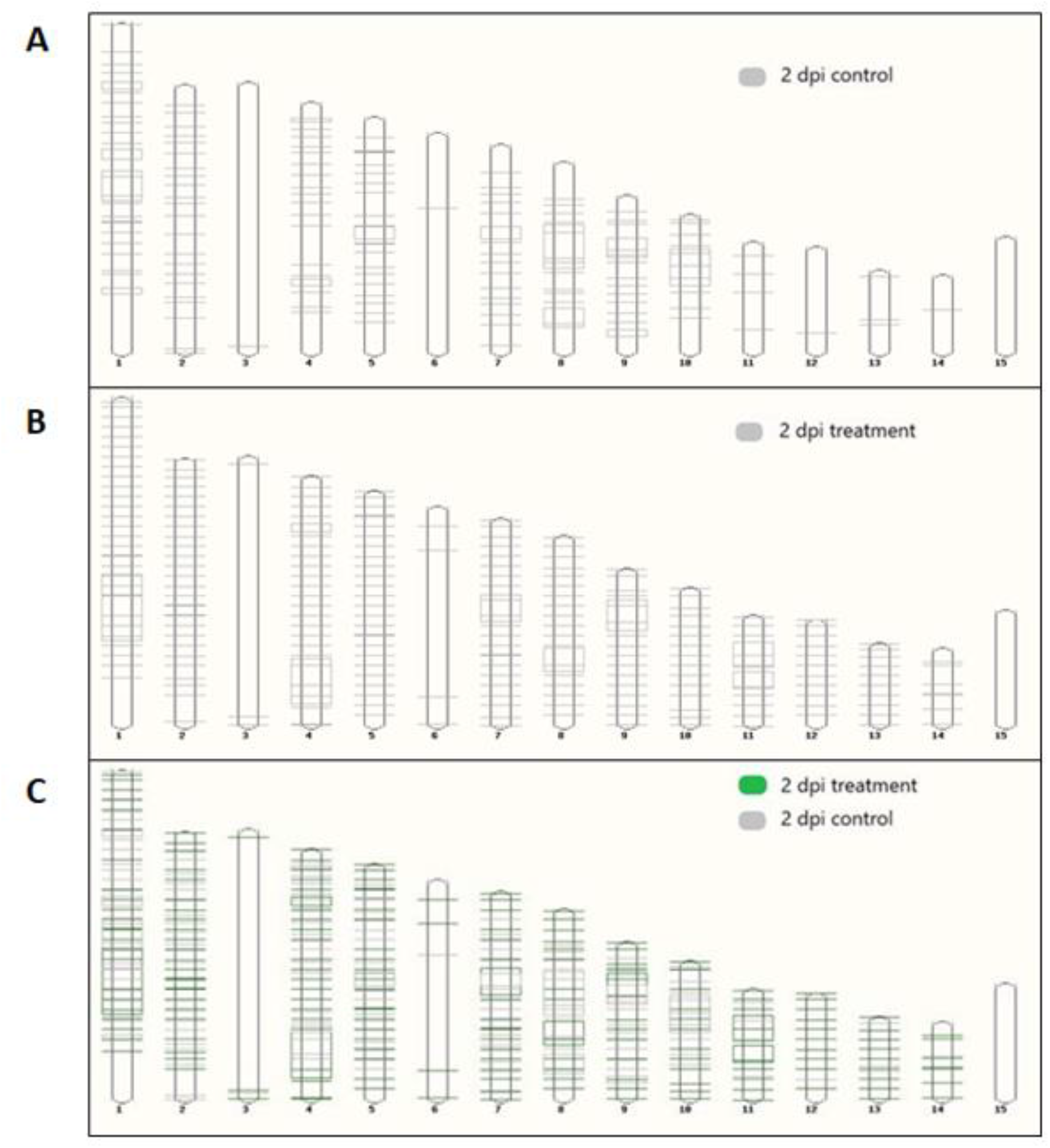
Fol 4287 karyotype with alignment of JAGH3 strain transcripts present during vanilla infection at 2 dpi. A) the transcripts corresponding to the control treatment. B) transcripts present in the treatment with the pathogenic strain JAGH3. C) intersection of the transcripts presents in the control treatment and the treatment with the JAGH3 strain.

In figure 2 we detected the transcripts corresponding to the libraries of the root treatment that were inoculated with the JAGH3 strain at 2 dpi and their location in the Fol 4287 reference genome.

**Figure 2.**
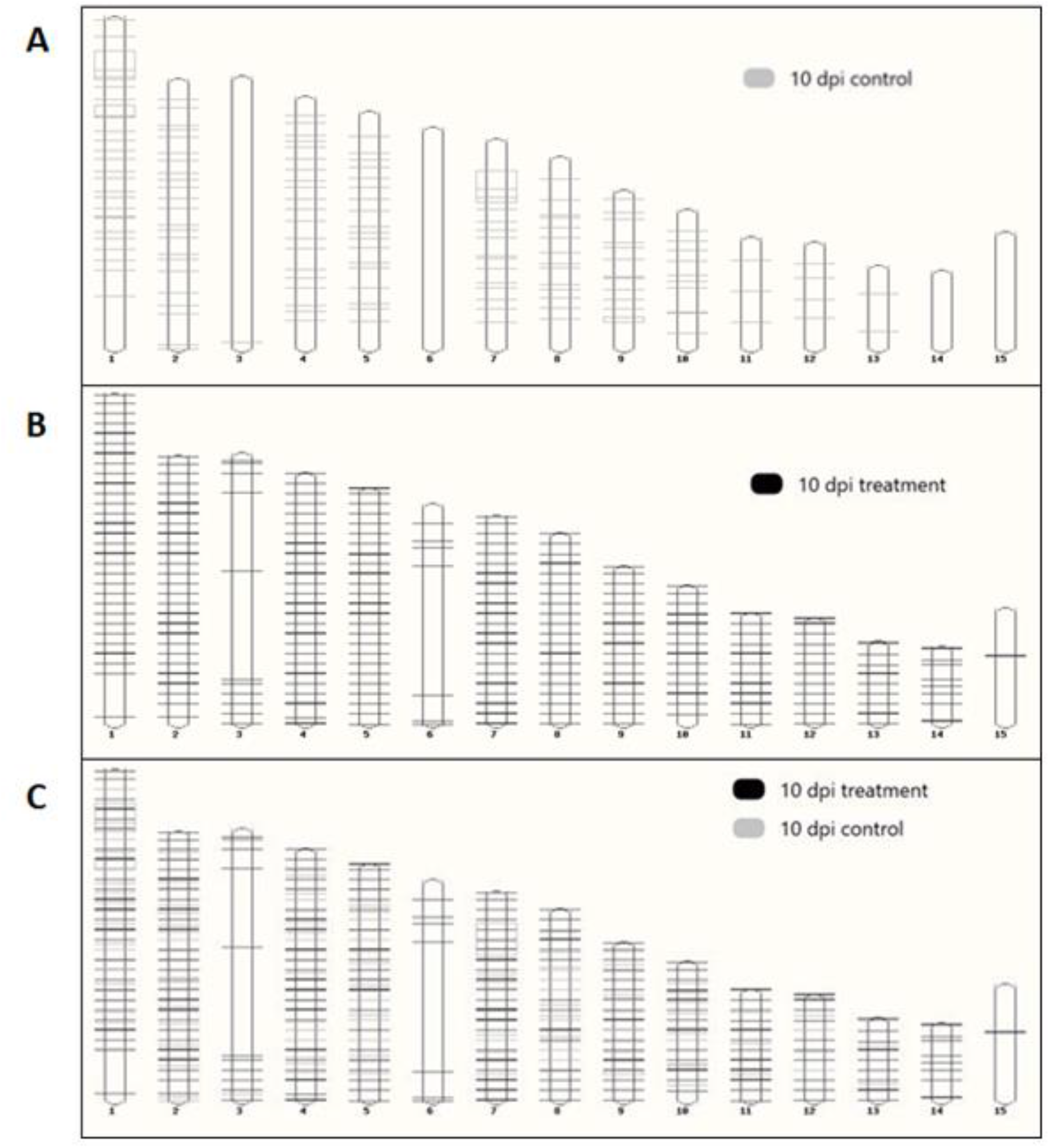
Fol 4287 karyotype with alignment of JAGH3 strain transcripts present during vanilla infection at 10 dpi. A) the transcripts corresponding to the control treatment. B) transcripts present in the treatment with the pathogenic strain JAGH3. C) intersection of the transcripts presents in the control treatment and the treatment with the JAGH3 strain.

Concerning the treatments where the JAGH3 strain was inoculated, in the Fol 4287 karyotype we found a greater number of transcripts present in each of the chromosomes, not only in the core genome but also in the supernumerary chromosomes. In fact, in the treatment at 2 dpi, in the core genome, we found the presence of transcripts in chromosomes 1, 2, 4, 5, 7, 8 and 9 with a clear increase in the number of transcripts present in each of these chromosomes against the control, being even more contrasting the increase in the number of transcripts observed in chromosomes 10, 11, 12, 13 and 14. The latter being considered as supernumeraries and associated with pathogenicity in *Fox*.

Finally, this increase in the number of transcripts present in the Fol 4287 karyotype observed in the treatment with the JAGH3 strain was evidenced in a contrasting way compared to the control treatment, at 10 dpi, where we found a significant increase on the core genome chromosomes and in the supernumerary chromosomes.

A conspicuous aspect that we must remark is the presence of components of pathogenicity islands in Supplementary Figure 1, present only in the libraries of the treatment with the JAGH3 strain at 10 dpi. Thus, showing the participation of this component in late infection and could be associated with the establishment of the necrotrophic behavior during the infection caused by *Fov* in vanilla.

### Metabolic pathways of the JAGH3 strain related to the infection process of the *V. planifolia* Jacks root

To elucidate the metabolic pathways in which the annotated *Fov* JAGH3 strain genes may be playing a crucial role during the infection in the vanilla root. We performed a functional prediction annotation and metabolic mapping using KEGG database (Kanehisa and Goto, 2000) (see supplementary material). Thus, the number of genes identified in the present work having a role in each of the pathways found was also obtained. In the supplementary table 1 and the supplementary table 2 there is a list of the metabolic pathways, and the number of genes associated with each pathway during early infection at 2 dpi (see Metabolic_routes_of_early_infection_at_2_dpi.xlsx) as well as late infection 10 dpi (see All_the_metabolic_routes_of_the_infection_dead_10_days.xlsx), respectively.

Interestingly, the number of genes corresponding to each of the metabolic pathways described above was considerably higher during late infection compared to early infection.

On the other hand, at 10 dpi we determined the presence of genes corresponding to the following metabolic pathways, which are only found during late infection: Penicillin and cephalosporin biosynthesis; Homologous recombination; Biotin metabolism; D-Arginine and D-ornithine metabolism; Atrazine degradation; Synthesis and degradation of ketone bodies; Sulfur relay system; C5-Branched dibasic acid metabolism; Polyketide sugar unit biosynthesis; Carotenoid biosynthesis; Selenocompound metabolism; Vitamin B6 metabolism; Carbapenem biosynthesis; Sesquiterpenoid and triterpenoid biosynthesis.

### Gene Ontology and KEGG Gene Enrichment analysis offer crucial insights about *Fov* pathogenesis in *V. planifolia* Jacks

Concerning the biological processes of GO analysis, we obtained 125 and 43 statistically significant at 2 dpi and 10 dpi, respectively. Where at 2 dpi (Figure 3) we observed terms involved in sporulation such as cell wall biogenesis and external encapsulating structure organization terms. Also, mechanisms possibly linked to virulence and infection as we found terms associated with protein, peptide, and organic substance transport. At 10 dpi (Figure 4) there were mostly regulatory mechanisms of nucleic acids, metabolic processes of macromolecules and organic substances and terms possibly related to sporulation including cell cycle. These data suggest that pathogenic mechanisms like sporulation, toxins and pathogenic effectors remain active along the infection process from early to late stages.

**Figure 3.**
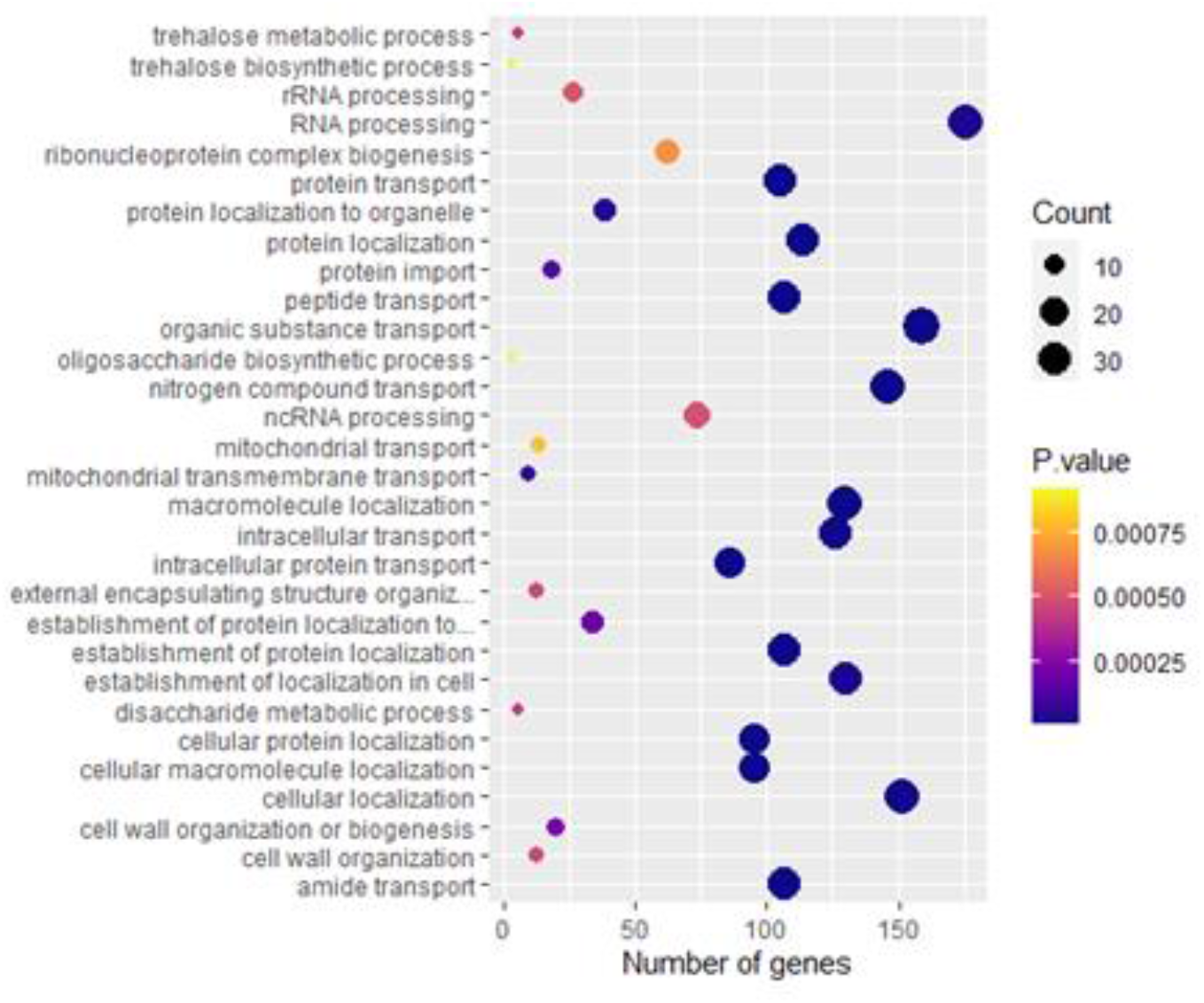
Gene Ontology of top 30 biological processes of Fov at 2 dpi in *V. planifolia* Jacks.

**Figure 4.**
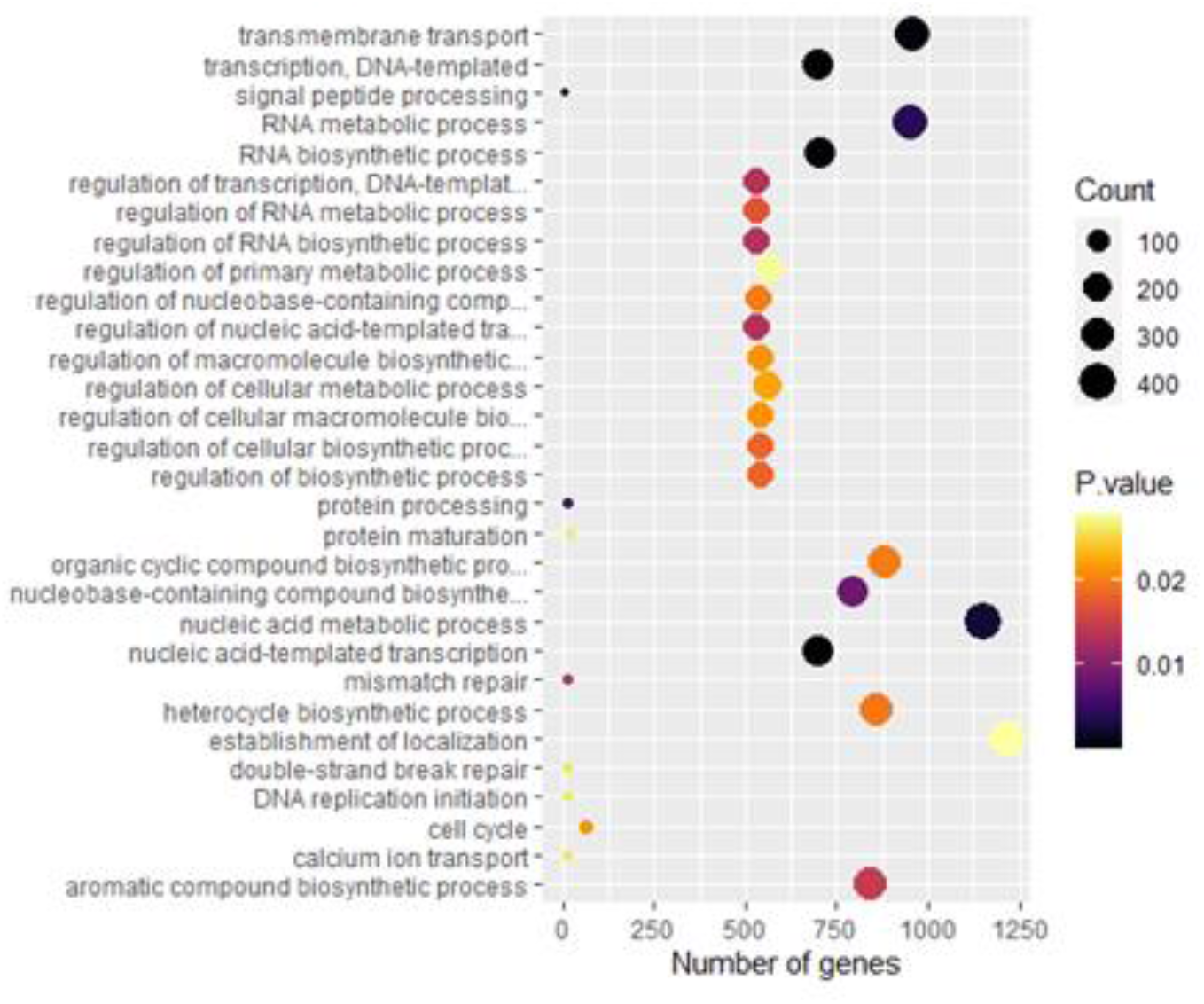
Gene Ontology of top 30 biological processes of Fov at 10 dpi in *V. planifolia* Jacks.

Regarding the metabolism, we found 21 and 46 statistically significant metabolic pathways at 2 dpi and 10 dpi, respectively. At 2 dpi (Figure 5) we observed pathways involved with reproduction and biosynthesis of metabolites and other biomolecules. In a similar manner we found pathways linked with these processes at 10 dpi (Figure 6). Additionally, there were other types of biomolecules synthesized at the late stage of infection. These results further confirm the previous found in GO analysis where biological processes and now metabolic pathways are associated with pathogenesis and stay operating along the infection, thus explaining the pathogenic behavior of *Fov*.

**Figure 5.**
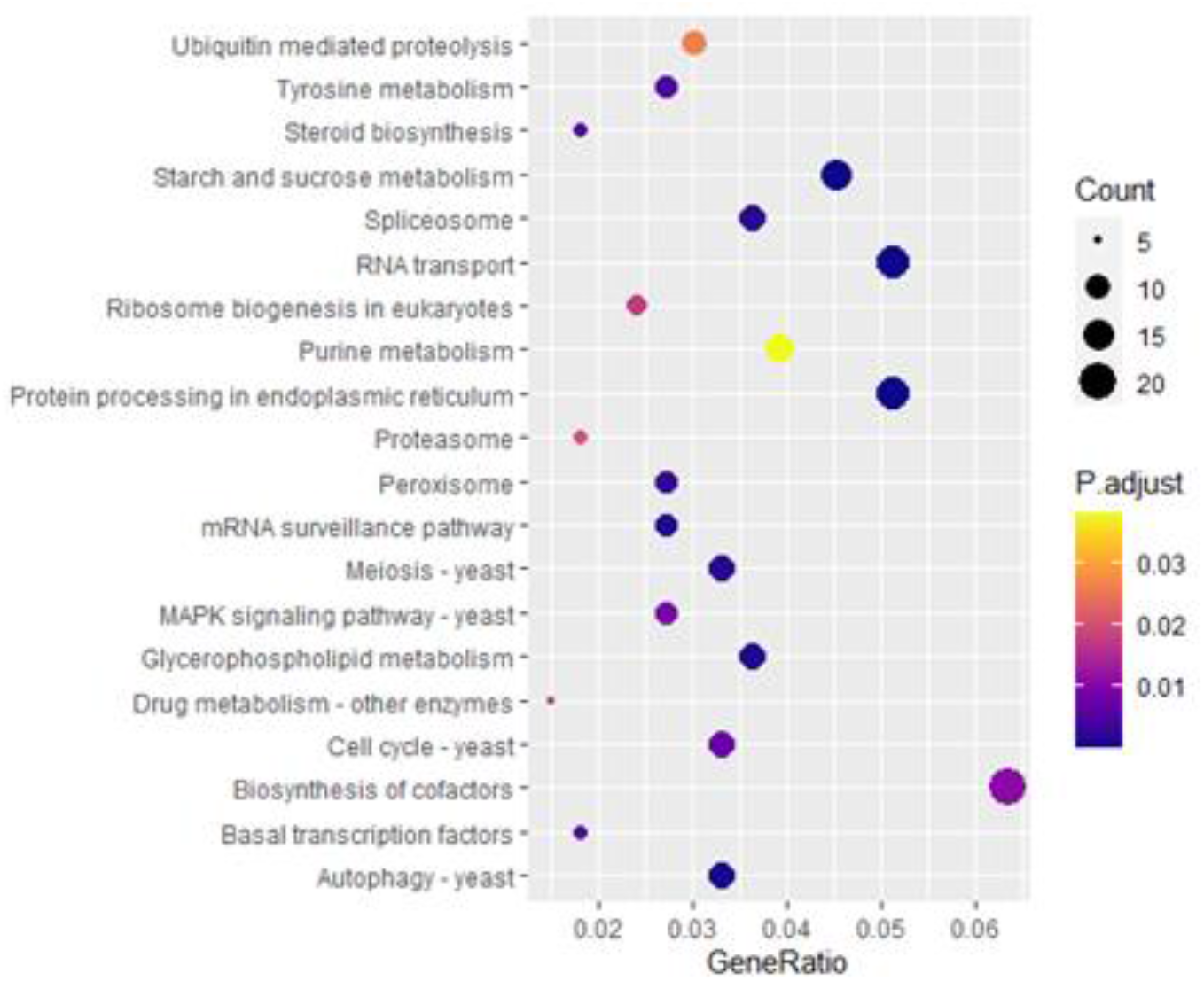
KEGG Gene Enrichment Analysis of top 20 metabolic pathways of Fov at 2 dpi in *V. planifolia* Jacks.

**Figure 6.**
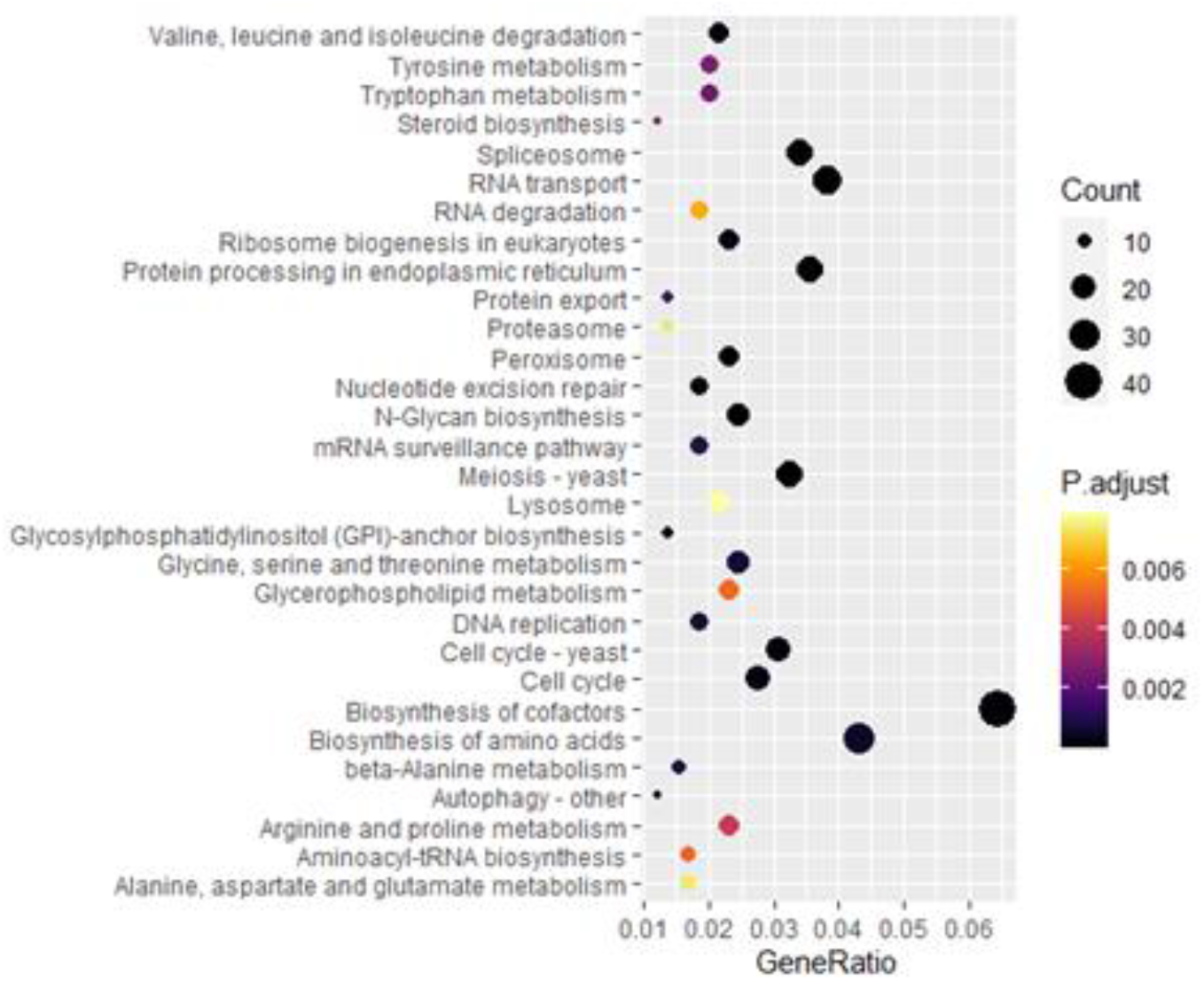
KEGG Gene Enrichment Analysis of top 30 metabolic pathways of Fov at 10 dpi in *V. planifolia* Jacks.

### *De novo* annotation as a functional prediction of *Fov* transcripts shed light on the role of pathogenesis related genes in the plant-host interaction

To characterize the molecular players that may play a main role during the pathogenesis in vanilla we did a *de novo* annotation as a functional prediction of *Fov* genes during these processes, firstly categorizing the shared and not shared genes between these two-time frames (Table 2).

**Table 2.** Shared and not shared genes numbers between early and late stage of infection.

| Category | Gene Count |
| --- | --- |
| Early stage infection | 3045 |
| Late stage infection | 7734 |
| Exclusive of early stage infection | 1334 |
| Exclusive of late stage infection | 6023 |

Among the annotated genes, we found virulence and hypervirulence factors during both stages of infection, interestingly these virulence factors increase in number and types during the late stage of infection (Table 3) in contrast with the early stage of infection (Table 4), this could mean that different virulence factors participate during different time frames along the disease.

**Table 3.** Hypervirulence and virulence factors of Fov at 2 dpi in *V. planifolia* Jacks.

| Gene ID | Coding protein | Fusarium species |
| --- | --- | --- |
| **FOXG_15434** | Pectin lyase fold/virulence factor | F. oxysporum f. sp. vanillae |
| **FOXG_05948** | Pectin lyase fold/virulence factor | F. oxysporum f. sp. vanillae |
| **FOXG_09893** | Pectin lyase fold/virulence factor | F. oxysporum f. sp. vanillae |
| **FOXG_09744** | Pectin lyase fold/virulence factor | F. oxysporum f. sp. vanillae |
| **FOXG_13795** | Pectin lyase fold/virulence factor | F. oxysporum f. sp. vanillae |
| **FOXG_08862** | Pectin lyase fold/virulence factor | F. oxysporum f. sp. vanillae |
| **FOXG_12264** | Pectin lyase fold/virulence factor | F. oxysporum f. sp. vanillae |
| **FOXG_02513** | Pectin lyase fold/virulence factor | F. oxysporum f. sp. vanillae |
| **FOXG_13794** | Pectin lyase fold/virulence factor | F. oxysporum f. sp. vanillae |
| **FOXG_04765** | Pectin lyase fold/virulence factor | F. oxysporum f. sp. vanillae |
| **FOXG_11739** | Pectin lyase fold/virulence factor | F. oxysporum f. sp. vanillae |
| **FOXG_14695** | Pectin lyase fold/virulence factor | F. oxysporum f. sp. vanillae |
| **FOXG_13051** | Pectin lyase fold/virulence factor | F. oxysporum f. sp. vanillae |
| **FOXG_12330** | Pectin lyase fold/virulence factor | F. oxysporum f. sp. vanillae |
| **FOXG_12863** | Hypervirulence associated protein, TUDOR domain | F. oxysporum f. sp. vanillae |
| **FOXG_15415** | Pectin lyase fold/virulence factor | F. oxysporum f. sp. vanillae |
| **FOXG_09894** | Pectin lyase fold/virulence factor | F. oxysporum f. sp. vanillae |

**Table 4.** Hypervirulence and virulence factors of Fov at 10 dpi in *V. planifolia* Jacks.

| Gene ID | Coding protein | Fusarium species |
| --- | --- | --- |
| FOXG_17741 | Pectin lyase fold/virulence factor | *F. oxysporum* f. sp. *vanillae* |
| FOXG_15434 | Pectin lyase fold/virulence factor | *F. oxysporum* f. sp. *vanillae* |
| FOXG_05948 | Pectin lyase fold/virulence factor | *F. oxysporum* f. sp. *vanillae* |
| FOXG_05861 | Egh16-like virulence factor | *F. oxysporum* f. sp. *vanillae* |
| FOXG_13331 | Pectin lyase fold/virulence factor | *F. oxysporum* f. sp. *vanillae* |
| FOXG_09893 | Pectin lyase fold/virulence factor | *F. oxysporum* f. sp. *vanillae* |
| FOXG_11096 | Pectin lyase fold/virulence factor | *F. oxysporum* f. sp. *vanillae* |
| FOXG_13795 | Pectin lyase fold/virulence factor | *F. oxysporum* f. sp. *vanillae* |
| FOXG_08862 | Pectin lyase fold/virulence factor | *F. oxysporum* f. sp. *vanillae* |
| FOXG_12264 | Pectin lyase fold/virulence factor | *F. oxysporum* f. sp. *vanillae* |
| FOXG_02509 | Pectin lyase fold/virulence factor | *F. oxysporum* f. sp. *vanillae* |
| FOXG_13794 | Pectin lyase fold/virulence factor | *F. oxysporum* f. sp. *vanillae* |
| FOXG_11739 | Pectin lyase fold/virulence factor | *F. oxysporum* f. sp. *vanillae* |
| FOXG_05407 | Pectin lyase fold/virulence factor | *F. oxysporum* f. sp. *vanillae* |
| FOXG_16516 | Pectin lyase fold/virulence factor | *F. oxysporum* f. sp. *vanillae* |
| FOXG_13051 | Pectin lyase fold/virulence factor | *F. oxysporum* f. sp. *vanillae* |
| FOXG_13801 | Pectin lyase fold/virulence factor | *F. oxysporum* f. sp. *vanillae* |
| FOXG_06038 | Pectin lyase fold/virulence factor | *F. oxysporum* f. sp. *vanillae* |
| FOXG_12863 | Hypervirulence associated protein, TUDOR domain | *F. oxysporum* f. sp. *vanillae* |
| FOXG_10052 | Pectin lyase fold/virulence factor | *F. oxysporum* f. sp. *vanillae* |
| FOXG_15415 | Pectin lyase fold/virulence factor | *F. oxysporum* f. sp. *vanillae* |
| FOXG_11740 | Pectin lyase fold/virulence factor | *F. oxysporum* f. sp. *vanillae* |
| FOXG_14660 | Pectin lyase fold/virulence factor | *F. oxysporum* f. sp. *vanillae* |
| FOXG_13103 | Pectin lyase fold/virulence factor | *F. oxysporum* f. sp. *vanillae* |
| FOXG_09744 | Pectin lyase fold/virulence factor | *F. oxysporum* f. sp. *vanillae* |
| FOXG_02712 | Egh16-like virulence factor | *F. oxysporum* f. sp. *vanillae* |
| FOXG_02513 | Pectin lyase fold/virulence factor | *F. oxysporum* f. sp. *vanillae* |
| FOXG_16409 | Pectin lyase fold/virulence factor | *F. oxysporum* f. sp. *vanillae* |
| FOXG_14695 | Pectin lyase fold/virulence factor | *F. oxysporum* f. sp. *vanillae* |
| FOXG_10728 | Pectin lyase fold/virulence factor | *F. oxysporum* f. sp. *vanillae* |
| **FOXG_12330** | Pectin lyase fold/virulence factor | *F. oxysporum* f. sp. *vanillae* |
| **FOXG_10062** | Pectin lyase fold/virulence factor | *F. oxysporum* f. sp. *vanillae* |
| **FOXG_17000** | Pectin lyase fold/virulence factor | *F. oxysporum* f. sp. *vanillae* |
| **FOXG_10173** | Egh16-like virulence factor | *F. oxysporum* f. sp. *vanillae* |

Additionally, at 2 dpi we obtained genes associated with sporulation (Table 5), necrotic/pathogenic activity (Table 6) and fusaric acid (Table 7). These results further confirm the molecular mechanisms which establish the pathogenic toolbox used by *Fov* during early infection stages.

**Table 5.** Sporulation genes of Fov at 2 dpi in *V. planifolia* Jacks.

| Gene ID | Coding protein | Fusarium species |
| --- | --- | --- |
| **FOXG_09277** | Stage II sporulation protein E (SpoIIE) | *F. oxysporum* f. sp. *vanillae* |
| **FOXG_08691** | Sporulation-specific protein Spo7 | *F. oxysporum* f. sp. *vanillae* |

**Table 6.** Necrosis/pathogenic genes of Fov at 2 dpi in *V. planifolia* Jacks.

| Gene ID | Coding protein | Fusarium species |
| --- | --- | --- |
| **FOXG_09418** | Clp amino terminal domain, pathogenicity island component | *F. oxysporum* f. sp. *vanillae* |
| **FOXG_07723** | Necrosis inducing protein (NPP1) | *F. oxysporum* f. sp. *vanillae* |

**Table 7.** Fusaric acid related genes of Fov at 2 dpi in *V. planifolia* Jacks.

| Gene ID | Coding protein | Fusarium species |
| --- | --- | --- |
| **FOXG_09823** | Fusaric acid resistance protein-like | *F. oxysporum* f. sp. *vanillae* |
| **FOXG_08593** | Fusaric acid resistance protein-like | *F. oxysporum* f. sp. *vanillae* |
| **FOXG_16912** | Fusaric acid resistance protein-like | *F. oxysporum* f. sp. *vanillae* |

Meanwhile, at 10 dpi genes involved in pathogenic processes found at 2 dpi were observed. However, there were remarkable differences in comparison with the late stage of infection, where there was conidiation (Table 8), pathogen-effector (Table 9) and Fusarinine (Table 10) related genes that were absent during the early stage of infection. Therefore, suggesting that these pathogenic processes are late stage exclusive in *Fov-V. planifolia* Jacks pathosystem.

**Table 8.**
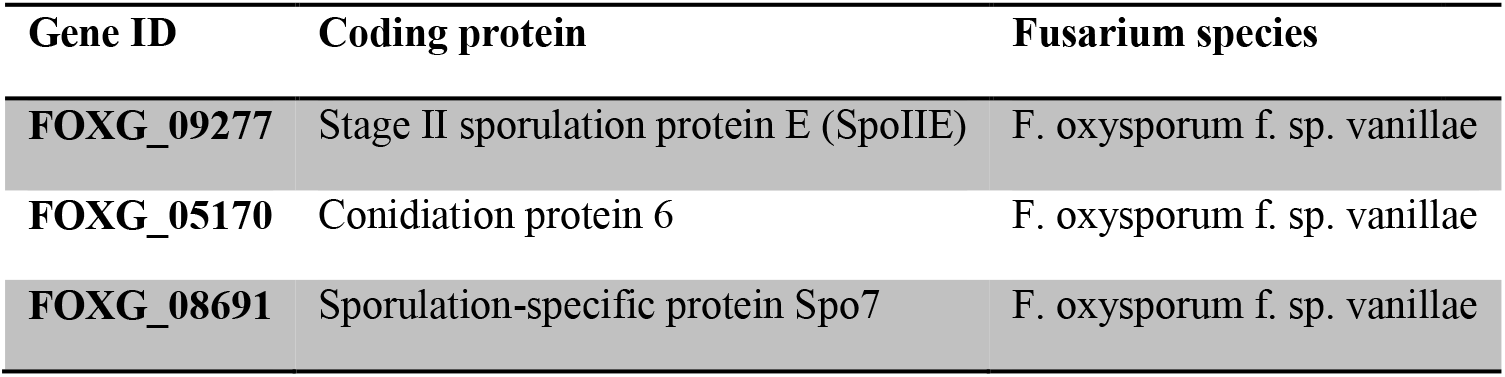
Conidiation and sporulation genes of Fov at 10 dpi in *V. planifolia* Jacks.

**Table 9.** Necrosis/pathogenic genes of Fov at 10 dpi in *V. planifolia* Jacks.

| Gene ID | Coding protein | Fusarium species |
| --- | --- | --- |
| **FOXG_06413** | Pathogen effector; putative necrosis-inducing factor | F. oxysporum f. sp. vanillae |
| **FOXG_09418** | Clp amino terminal domain, pathogenicity island component | F. oxysporum f. sp. vanillae |
| **FOXG_17014** | Necrosis inducing protein (NPP1) | F. oxysporum f. sp. vanillae |
| **FOXG_07723** | Necrosis inducing protein (NPP1) | F. oxysporum f. sp. vanillae |

**Table 10.**
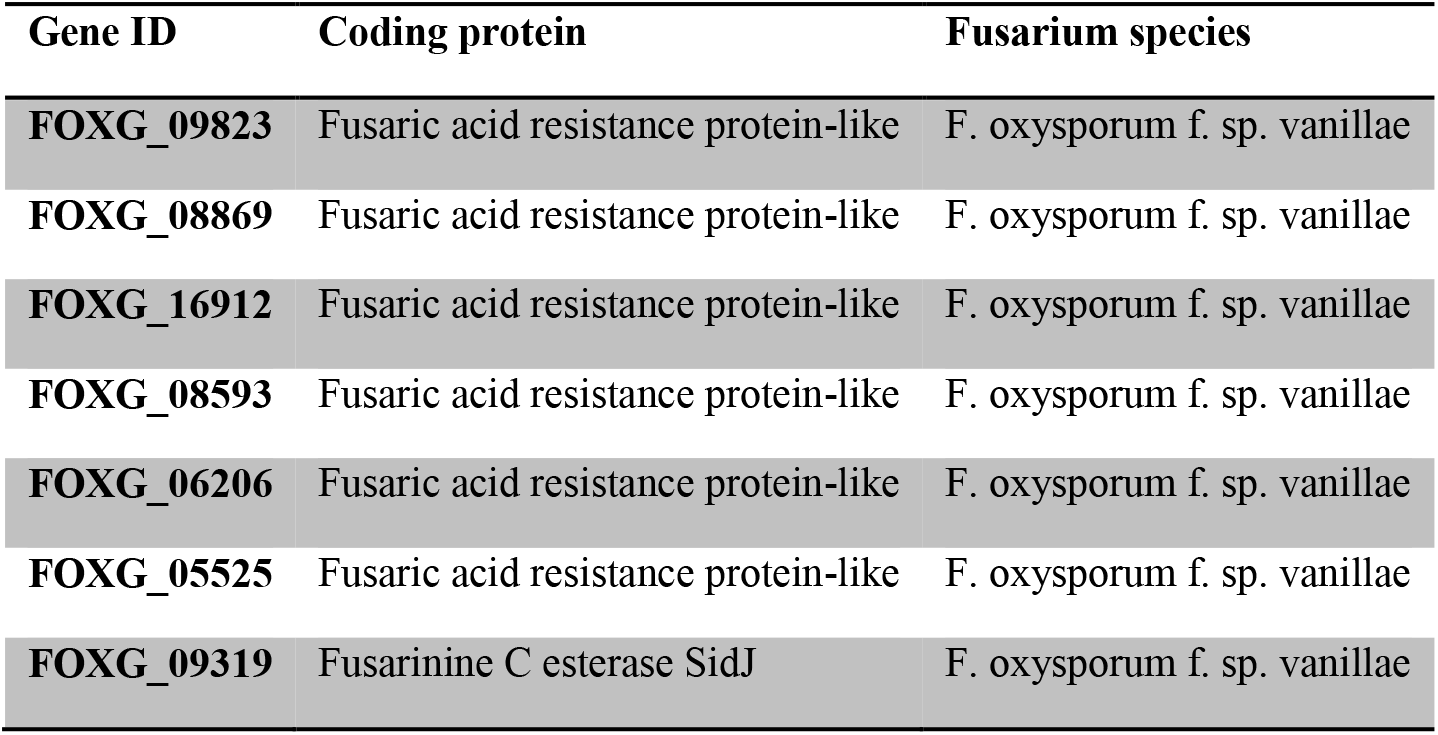
Fusarinine and fusaric acid related genes of Fov at 10 dpi in *V. planifolia* Jacks.

## Discussion

### Use of sugar as an energy source and the growth of conidia in early *Fox* infection in vanilla root

As previously described, the synthesis of carbohydrates, as well as the production of sugar, alcohol and organic acids is one of the most active metabolic pathways of *Fox* metabolism during the germination process of conidia (Sharma et al., 2016). This, being one of the first differences that we observed in this work in relation to the control treatment, where the pathogenic strain JAGH3 was not inoculated. The significant presence of genes corresponding to the synthesis of carbohydrates, production of alcohol and organic acids in the treatments with the JAGH3 strain in both time frames of infection was detected. We also showed the presence of transcripts related to carbohydrate metabolism, oxidative phosphorylation, glycolysis, and the pentose phosphate pathway as metabolic pathways related to the germination of *Fox* conidia (Sharma et al., 2016). In addition to the mentioned study, there is evidence that sugar, as an easily oxidized carbon source, supplies the demand for the conidia germination process (Cochrane et al., 1963; Griffin 1970a; and Griffin, 1970b). In fact, at 2 dpi we found the presence of genes of hexokinase, trehalose 6-phosphate synthase, beta-fructofuranosidase, neutral trehalase, murein transglycosylase, alpha-trehalose-phosphate synthase, trehalose 6-phosphate synthase, endoglucanase type B, beta-glucosidase, glucoamylase, alpha-amylase, glycosyl hydrolase family, corresponding to the metabolism of starch and sucrose (see supplementary material). The above is evidence of the use of an energy source during *Fov* infection in vanilla root, which previous works have associated with the growth of hyphae in *Fox* during the germination of conidia, a biological process that is associated with the early stages of infection.

At 10 dpi we found the following genes linked to the metabolism of starch and sucrose: beta-glucosidase, murein transglycosylase, glucan 1,6-alpha-glucosidase, beta-glucosidase G, trehalase, beta-glucosidase, alpha-glucosidase, endoglucanase, beta-glucosidase K, glycogen phosphorylase, endoglucanase, oligo-1,6-glucosidase, alpha alpha-trehalse, glucan 1,3-beta-glucosidase. In addition to the genes involved with glycolysis and gluconeogenesis: alcohol dehydrogenase (NADP +), aldehyde dehydrogenase (NAD +), pyruvate decarboxylase, aldose 1-epimerase, which prefer propanol. This shows the involvement of carbohydrate metabolism in the late phase of *Fov* infection in vanilla.

### Amino acid metabolism in *Fox* infection in vanilla root

The participation of amino acid metabolism in plant-pathogen interaction and its participation in the establishment of pathogenicity has been previously documented (Phillips et al., 2004; Jonkers et al., 2009; Scandiani et al., 2014; Tchameni et al., 2012; Okada and Matsubara, 2021; Kasote et al., 2020). At 2 dpi, different genes related to amino acid biosynthesis were determined, such as: aspartate kinase; pyruvate carboxylase; tagatose 1,6-diphosphate aldolase; asparagine synthetase; ribose-phosphate pyrophosphokinase 3, 6-phosphofructokinase, acetylornithine aminotransferase, mitochondrial; tryptophan synthase and branched-chain amino acid aminotransferase. Jastrzębowska and Gabriel in 2015, proved that the inhibition of amino acid biosynthesis, by means of chemical substances that act as antifungals (N-(5-substituted-1,3,4-thiadiazol-2-yl) cyclo-propanecarboxamides), significantly reduces *Fox* growth in minimal medium. Likewise, these authors refer that auxotrophic mutants of human pathogenic fungi impaired in biosynthesis of amino acids exhibit growth defects or at least reduced virulence under *in vivo* conditions (Jastrzębowska and Gabriel, 2015). At 10 dpi, we also confirmed the participation of genes associated to amino acid biosynthesis in the growth of conidia, by determining the presence of genes corresponding to: cystathionine gamma-lyase; anthranilate phosphoribosyltransferase; cysteine synthase A; mitochondrial homocitrate synthase; argininosuccinate lyase; dTDP-4-dehydrorhamnose reductase; 3-isopropylmalate dehydrogenase; dihydroxy-acid dehydratase, mitochondrial; ribose 5-phosphate isomerase B; serine hydroxymethyltransferase, mitochondrial; homoserine dehydrogenase; threonine aldolase; imidazoleglycerol-phosphate dehydratase; argininosuccinate synthase; acetylornithine deacetylase; dihydrodipicolinate synthase; glutamine synthetase; homoisocitrate dehydrogenase; pyrroline-5-carboxylate reductase; tryptophan synthase, beta subunit; 3-dehydroquinate synthase; L-2-aminoadipate reductase; glycine hydroxymethyltransferase; S-adenosylmethionine synthetase; 3-deoxy-7-phosphoheptulonate synthase; threonine dehydratase.

Also, at 2 dpi we found the presence of genes linked to the synthesis of tyrosine such as: acylpyruvate hydrolase, primary-amine oxidase, aldehyde dehydrogenase (NAD +), tyrosinase, gentisate 1,2-dioxygenase, 4-hydroxyphenylpyruvate dioxygenase, primary - amine oxidase, homogentisate 1,2-dioxygenase, and fumarylacetoacetase. Whose presence and importance in the genomic context were reported in filamentous fungi Greene et al., (2014). For its part, at 10 dpi, we detected the following genes involved with the tyrosine metabolism pathway: catechol O-methyltransferase, succinate-semialdehyde dehydrogenase (NADP +); maleylacetoacetate isomerase; copper amine oxidase; succinate-semialdehyde dehydrogenase / glutarate-semialdehyde dehydrogenase; monoamine oxidase; 1,2-dioxygenase homogentisate; alcohol dehydrogenase, propanol-preferring; and aspartate aminotransferase, cytoplasmic.

At 10 dpi we observed transcripts related to arginine and proline metabolism: glutamate 5-kinase; 1-pyrroline-5-carboxylate dehydrogenase; ornithine aminotransferase; 1-pyrroline-5-carboxylate dehydrogenase; N1-acetylpolyamine oxidase; N1-acetylpolyamine oxidase; aldehyde dehydrogenase (NAD +); D-amino-acid oxidase; monoamine oxidase; proC; 1-pyrroline-5-carboxylate dehydrogenase; amidase; ornithine decarboxylase; agmatine deiminase; polyamine oxidase; aspartate aminotransferase, cytoplasmic; 4-hydroxy-2-oxoglutarate aldolase. Boedi et al in 2016 reported the comparison by transcriptome analysis of the pathogenic (biotrophic) phase against the saprophytic phase in *F. graminearum* (Boedi et al., 2016). Finding in pathogenesis an increase in the order of at least two times in magnitude in the expression of genes corresponding to the arginine and proline metabolism pathway. Indicating that the metabolism of arginine and proline plays an important role in the growth of hyphae, participating in the biotrophic phase of *Fox* in plants.

### Protein metabolism in *Fox* infection in vanilla root

Concerning the protein metabolism its participation in the establishment of pathogenicity has been widely reported (Kumar et al., 2016; Kong et al., 2019). We found at 2 dpi the presence of genes associated with the processing of proteins in the endoplasmic reticulum, such as: nuclear protein localization protein 4, oligosaccharyl transferase complex subunit OST4, protein disulfide-isomerase erp38, translocation protein SEC63, protein transporter SEC23, mannosyl-oligosaccharide alpha-1,2-mannosidase, hypothetical protein, hypothetical protein, protein disulfide-isomerase, Cullin 1, hypothetical protein, hypothetical protein, translation initiation factor 2 subunit 1, IRE protein kinase, DnaJ like subfamily B member 12, protein transporter SEC61 subunit beta, protein OS-9. Wang et al., (2020) report that the use of myriocin as an antifungal negatively affects the stability of the membrane of *F. oxysporum* f. sp. *niveum*. When analyzing the combined analysis between the transcriptome and proteome revealed that the expression of some membrane-related genes and proteins, mainly those associated with sphingolipid metabolism, glycerophospholipid metabolism, steroid biosynthesis, ABC transporters and protein processing in the endoplasmic reticulum, was disordered.

Also, in the treatment of 2 dpi we detected the presence of genes linked to proteolysis mediated by ubiquitination, these genes correspond to: F-box and WD-40 domain-containing protein MET30, F-box and WD-40 domain-containing protein CDC4, ubiquitin-conjugating enzyme (huntingtin interacting protein 2), SUMO-conjugating enzyme ubc9, Cullin 1, ubiquitin-like 1-activating enzyme E1 B, ubiquitin-conjugating enzyme E2 R and pre-mRNA-processing factor 19. Recently, numerous studies revealed that F-box proteins are required for fungal pathogenicity (Liu and Xue, 2011). Proteome analysis in *F. oxysporum* showed the role of Fb1 protein in proteasome degradation and susceptibility as potential targets of ubiquitination (Manikandan et al., 2018; Miguel-Rojas and Hera, 2013). A proteomic approach in *F. oxysporum* f.sp. *lycopersici* was done by Miguel-Rojas and Hera (2013) where they identified the differential pattern of proteins involved in vesicle blocking and proteasome degradation (Manikandan et al., 2018). The function of F-box proteins and their potential as part of the ubiquitin-proteasome system (UPS) has been reported in several fungal pathogens, including those that cause human infections (such as *C. neoformans* and *C. albicans*) or plant diseases (such as *Fusarium* species and *M. oryzae*) (Liu and Xue, 2011). Fungi such as *Fusarium* species contain much larger numbers of F-box proteins than yeast, approximately 60~94 in *Fusarium* species compared to around 20 in yeasts (*S. cerevisiae*, *C. albicans*, or *C. neoformans*). Suggesting that F-box proteins contribute to the regulation of a more complex developmental and metabolic processes occurring in filamentous fungi (Liu and Xue, 2011).

### Autophagy in *Fox* infection in vanilla root

Autophagy has been described as one of the main routes of cell traffic and recycling, also playing a relevant role in pathogenesis (Lv et al., 2017). These authors, in 2017 reported genome-wide identification and characterization of autophagy-related genes (ATGs) in the wheat pathogenic fungus *F. graminearum* identifying 28 genes associated with the regulation and operation of autophagy; also finding that subsets of autophagy genes were necessary for asexual / sexual differentiation and deoxynivalenol (DON) production, respectively. Finally, they concluded that autophagy plays a critical function in growth, asexual / sexual sporulation, deoxynivalenol production and virulence in *F. graminearum*. At 2 dpi we found the presence of genes involved with the autophagy pathway such as: CAMK / CAMKL / AMPK protein kinase, hypothetical protein, synaptobrevin, serine / threonine-protein phosphatase PP2A catalytic subunit, Ras-like protein, translation initiation factor 2 subunit 1, CAMKK protein kinase. Kalhid et al., in 2019, found that autophagy assumes an imperative job in affecting the morphology, development, improvement and pathogenicity of *Fox*, through the study of an ATG gene that regulates its pathogenicity. Corral-Ramos et al., in 2015 demonstrated that autophagy mediates nuclear degradation after hyphal fusion and possesses a general function in the control of nuclear distribution in *Fox*; in a process regulated by ATG genes; during the vegetative growth of hyphae.

### Glycerophospholipid in *Fox* infection in vanilla root

In the present work we found the presence at 2 dpi of genes related to the glycerophospholipid metabolism pathway, as widely reported previously (van Aarle and Olsson, 2003; Bhandari et al., 2018; Zhang et al., 2020), such as: ethanolamine kinase, acetyltransferase, phosphatidylserine decarboxylase, phosphatidylserine decarboxylase, lysophospholipase NTE1, CDP-diacylglycerol-inositol 3-phosphatidyltransferase, ethanolamine-phosphate cytidylyltransferase, and NTE family protein. Joshi and Mathur in 1987 mentioned that *Fox* synthesizes up to 26.4% lipids in minimal medium, being glycerophospholipids and triacylglycerols, the latter predominating. The main glycerophospholipid is phosphatidylcholine, the others are phosphatidylethanolamine and phosphatidylinositol. Wang et al, reported in 2020, that Bacillus amyloliquefaciens LZN01 Myrozin can inhibit the growth of *F. oxysporum* f. sp. *leveum*. In a combined proteome and transcriptome analysis they found that it had expression of some membrane-related genes and proteins, mainly those related to sphingolipid metabolism, glycerophospholipid metabolism, steroid biosynthesis, ABC transporters and protein processing in the endoplasmic reticulum, was disordered. They concluded that myriocin has a significant antifungal activity due to its ability to induce membrane damage in this fungus. Zuo et al, in 2017 reported that sterol and glycerophospholipids are important cellular or subcellular membrane components and are required for cell growth and proliferation, maintenance of organelle morphology, signal transduction, and lipid homeostasis (Kihara and Igarashi, 2004; Nebauer et al., 2004). Furthermore, said authors reported that exposure to DT from Foc TR4 regulates the metabolism of steroids and glycerophospholipids, affecting the integrity of the cell membrane of FOC TR4 causing DT toxicity (Zuo et al., 2017). In late infection at 10 dpi, we found the following genes associated with the synthesis of Glycerophospholipid acetyltransferase, choline-phosphate cytidylyltransferase, phosphatidylserine decarboxylase, phosphatidylserine synthase, phosphatidylethansolipid synthey N-phosphatidylettransferase 3-methylhalserine-phosphate-synthase-3, phosphatidylethansinehydrogenase-3, phosphatidylethansinehydro-synthase-3 - 3-phosphate dehydrogenase, phospholipase A2, triacylglycerol lipase, phosphatidylserine decarboxylase, glycerol-3-phosphate dehydrogenase, ethanolamine kinase, acetylcholinesterase, phospholipase D1 / 2, phosphatidylaglycerine decarboxylasecerine A2, phosphatidylaglysecerine decarboxysecerine A2, phosphatidylaglysecerine decarboxysecerine A2, phosphatidylaglysecerine decarboxysecerine 1 - serine O-phosphatidyltransferase, diacylglycerol diphosphate phosphatase / phosphatidate phosphatase.

### Bioinformatic analysis results throw light on the underlying molecular mechanisms used in pathogenicity in the *Fov-V. planifolia* Jacks pathosystem

In FOSC, the infection has been described in *F. oxysporum* f. sp. *lycopersici*. This pathogenic process is divided into four main events. It starts with the hyphae adhesion to the plant surface, continuing with the penetration of the plant structure which depends on biotic factors and environmental cues, considering that *F. oxysporum* is an opportunistic pathogen it needs an injury to enter (Bani et al., 2018). Once the fungus has established itself in the plant, the colonization process will begin, there mycelium and spores advance towards xylem vessels causing conidiation to start, later the spores and conidia are transport with the sap via xylem vessels, so these spores germinate and develop into conidiophores and conidia going through the perforation plates. Finally, the pathogen causes the occlusion of xylem vessels and hydric stress by the accumulation of mycelium and toxins, this being reflected in a variety of symptoms including: vascular wilt, abscission, chlorosis, and necrosis. Therefore, the death of the plant. (MacHardy and Beckman, 1981; Bishop and Cooper, 1983; Beckman, 1989; Olivain and Alabouvette, 1997; Lucas, 1998; Olivain and Alabouvette, 1999; Schneider and Carlquist, 2004; Groenewald, 2006; Olivain et al., 2006).

About the description of the infection in vanilla, there exists few reports of *F. oxysporum* f. sp. *radicis-vanillae* affecting this orchid. The symptoms start with the browning and death of underground and aerial roots, continuing with wrinkles in leaves and stem and ending with the death of the plant. As for the infection timing, it requires two days to achieve hyphae adhesion and eight days to complete the colonization phase (Koyyappurath et al., 2015; Koyyappurath et al., 2016).

In concurrence with the findings in *F. oxysporum* f. sp. *radicis-vanillae*, where they documented the germination of conidia and subsequent triggering of the infection at 2 dpi and the colonization of the root later frames of the disease (Koyyappurath et al., 2015). In this study we report, also at 2 dpi the presence of genes related to sporulation, fusaric acid, necrosis, virulence and hypervirulence factors. Thus, explaining the beginning of conidia germination and the genesis of the infection as these other pathogenesis-related genes act as key players during this early plant-host interaction. Also, in agreement, in the advanced stage of infection at 7 dpi and 9 dpi there was progression of the plant colonization and softening of the root tissue as consequence of the fusarium wilt. These symptoms could be justified with the data shown at 10 dpi, here we found genes related to conidiation, pathogenic effectors and fusarinine in addition to those reported at 2 dpi. These are foremostly linked to pathogenesis and tissular death. Therefore, exhibiting a predominant necrotic behavior as reported in this forma specialis (Koyyappurath et al., 2015; Koyyappurath et al., 2016). As these results indicate the possible pathogenic strategy of Fov we summarized it into a schematic representation (Figure 7).

**Figure 7.**
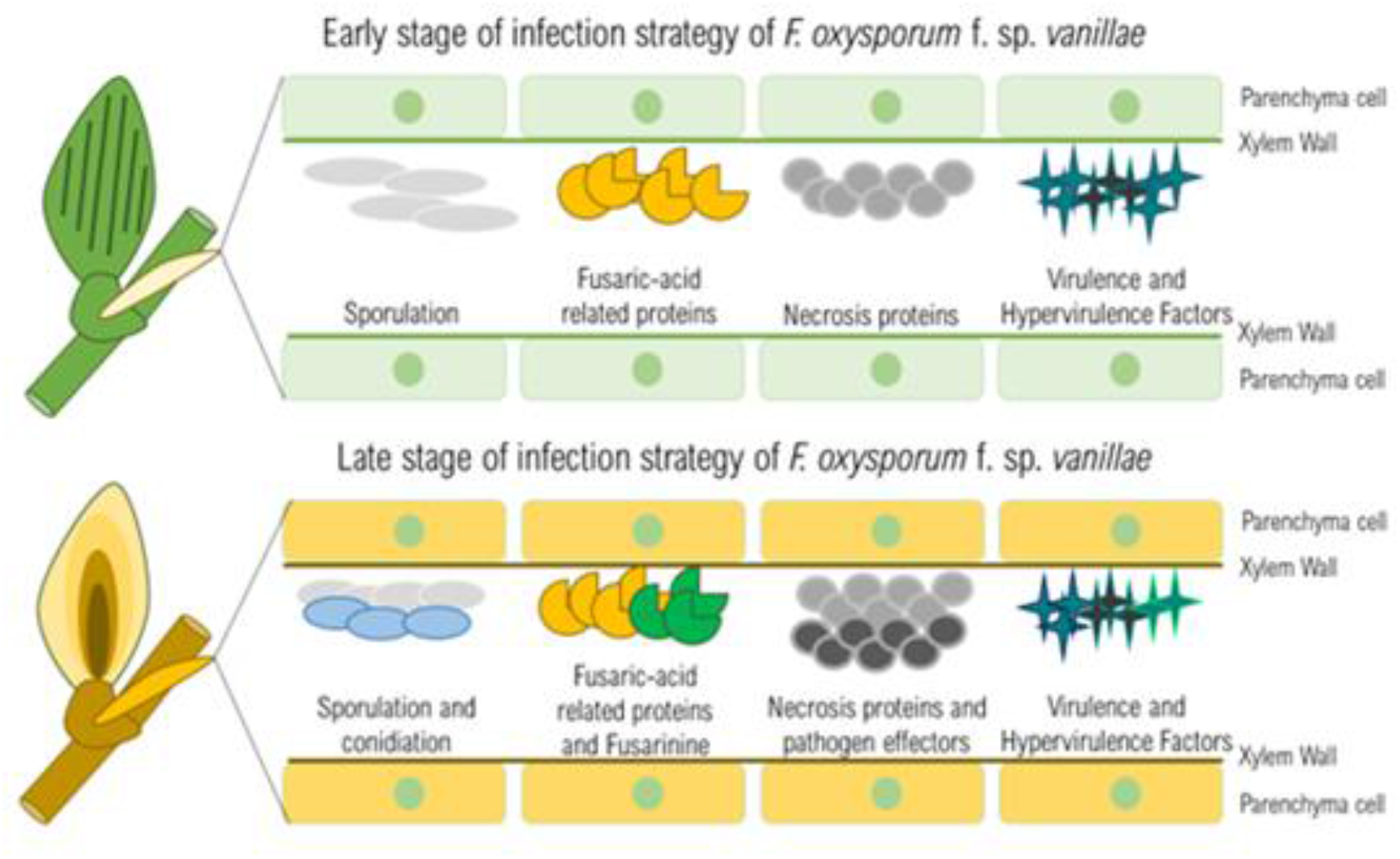
Hypothetical model of the Fov pathogenesis.

As growing evidence supports the hypothesis that pathogenicity of FOSC relies on transferable extra chromosomes and genes coding for Secreted in Xylem proteins also known as SIX (Ma et al., 2013). It was shown in this study the presence of genes belonging to pathogenicity chromosomes and pathogenicity islands as this hypothesis suggests. Nonetheless, we did not report the presence of SIX genes, therefore despite the similarities in the pathogenic strategy with other formae speciales, the findings in this work suggest that *F. oxysporum* f. sp. *vanillae* pathogenic capacity is SIX genes-independent but another pathogenic elements-dependent such as the previously described.

## Conclusions

The bioinformatic analysis of *Fov* transcripts during the infection in *V. planifolia* Jacks revealed that this forma specialis make use different pathogenesis associated genes along this process, among them we must remark sporulation, conidiation, fusaric acid, necrosis, pathogenic and virulence effectors as main molecular players. Events that are associated with primary metabolism, carbohydrates, amino acid metabolism, autophagy, and secondary metabolism. Hence, we propose that these allow *F. oxysporum* f. sp. *vanillae* to infect vanilla in a SIX genes-independent manner. Also, the presence of them could be explained by the pathogenicity islands and the genomic regions associated with supernumerary chromosomes found in *Fov*. Both act as carriers of genes involved with pathogenic activity. The results shown here highlight the key players that confer the capacity to infect vanilla to the deadliest pathogen affecting this crop.

## Methods

### Experimental framework

The experimental framework, that included the plant material collection, infectivity assays, total RNA extraction and the generation with the sequencing of cDNA libraries were done in our previous work in Solano-De la Cruz et al., 2019. The generated data was submitted to the GEO platform of NCBI-GenBank (Accession number: GSE134155).

### RNA-seq data processing: cleaning, alignment, and filtering

Quality of reads obtained from the high-throughput sequences was carried out using FASTQC version 0.11.2 (Andrews, 2010) and MultiQC version 1.0 (Ewels et al., 2016) software. Reads above 32 nt, without the presence of adapters were considered for further analysis. First, to filter and discard reads corresponding to the plant host, genome index and alignment of reads was done utilizing the STAR program version 2.7.2 (Dobin et al., 2012) using the reference genome of *F. oxysporum* f. sp. *lycopersici* strain Fol 4287 available on Ensembl Fungi and visualize the alignment output with MultiQC version 1.0 software. To further filter them based on alignment quality, Samtools version 1.9 (Li et al., 2009) and MultiQC for post-filtering visualization, were used.

### Genes identification and visualization of aligned genomic regions of Fov to Fol 4287 karyotype

To be able to identify the Fov transcripts participating in the infection, the GenIDs obtention was accomplished. To accomplish this, the intersection of the genomic coordinates of the filtered alignment results against the *F*. *oxysporum* f. sp. *lycopersici* strain Fol 4287 genome annotation was done using Bedtools version 2.25.0 (Quinlan and Hall, 2010), with the extraction of GenIDs lists using cut command and AWK, was done. On the other hand, the bed files with the genomic coordinates were used to map and visualize the Fov found genes in the *F*. *oxysporum* f. sp. *lycopersici* strain Fol 4287 karyotype using the visualization tool from Ensembl Fungi of Ensembl Genomes consortium (Howe et al., 2020) (available at https://fungi.ensembl.org/Fusarium_oxysporum/Location/Genome).

### Metabolic description of identified genes: annotation and mapping

To elucidate the metabolic pathways where the identified genes could be involved, an annotation and mapping of these biochemical processes were conducted. First, extracting the protein sequences of the identified genes using the Biomart-Ensembl Fungi database (Howe et al., 2020) to annotate them with the BlastKoala tool from KEGG platform (Kanehisa and Goto, 2000) (available at https://www.kegg.jp/kegg/tool/annotate_sequence.html). Later, mapping the annotation against the species-specific KEGG pathway maps belonging to *F. oxysporum* with the Search Pathway tool from KEGG platform (available at https://www.genome.jp/kegg/tool/map_pathway1.html).

### Gene Ontology Enrichment Analysis

GO enrichment analysis was executed using the topGO R package (Alexa et al., 2006), we applied the following criteria: p-value < 0.05 threshold, algorithm “classic” and “fisher” statistic. We utilized as query the obtained GeneIDs lists from *F. oxysporum* f. sp. *lycopersici* strain Fol 4287 in the Biomart-Ensembl Fungi database (Howe et al., 2020). Finally, we visualized this data via RStudio version 4.0.1 and ggplot2 package version 3.3.2 (Wickham, 2016).

### KEGG Gene Enrichment Analysis

We did the KEGG enrichment analysis using RStudio and the clusterProfiler package (Yu et al., 2012). To use the enrichKEGG function in this package, we used as input the KO identifiers obtained from BlastKoala annotation and filtered our results by applying adjusted p-value cut-offs < 0.05. For the visualization we used RStudio version 4.0.1 and ggplot2 R package version 3.3.2 (Wickham, 2016).

### *De novo* annotation of Fov identified genes

To decipher the potential functions of Fov gene transcripts found from out alignment against the Fol 4287 strain genome. We utilized the software InterproScan 5 version 5.41 (Jones et al., 2014). Which allows protein-coding sequences function prediction, using as reference databases Pfam and SUPERFAMILY.

## Supporting information

2_dpi_treatment_genes_list

10_dpi_treatment_genes_list

All_the_metabolic_routes_of_the_infection_dead_10_days

Metabolic_routes_of_early_infection_at_2_dpi

## Abbreviations

FOSC: *Fox* species complex
HGT: horizontal gene transfer
RSR: root and stem rot
Fov: *F. oxysporum* f. sp. *vanillae*
*Fox*: *F. oxysporum*
*Fol*: *F. oxysporum* f. sp. *lycopersici*
dpi: days post-inoculation
DON: deoxynivalenol.

## Acknowledgments

We are grateful to Dr. Carlos Lobato Tapia and Dr. Rosalinda Tapia López for reviewing this manuscript. To the University Massive Sequencing Unit of the Institute of Biotechnology, UNAM. To M.C Jerome Jean Verleyen for his help in the use of the Teopanzolco bioinformatics cluster. To the National Council of Science and Technology (CONACYT), for the scholarship 720145 granted to the first author EEEH to carry out his postgraduate studies. To the National Laboratory of Genomics for Biodiversity (LANGEBIO), CINVESTAV for the MAZORKA bioinformatic cluster services.

## Author’s contributions

MTSC, EEEH, MLR coordinated the study. MTSC, EEEH and MLR conceived and designed the experiments. MTSC and EEEH performed the bioinformatic analysis. MTSC and EEEH interpreted the results of the analyses. MTSC, EEEH, RPRZ, JAAG, JAG and MLR conceived and organized the manuscript structure. All authors contributed during the manuscript preparation and approved the final manuscript.

## Availability of data and materials

The datasets generated and/or analyzed during the current study are available in the GEO repository with accession number GSE134155 (https://www.ncbi.nlm.nih.gov/geo/query/acc.cgi?acc=GSE134155).

## Ethics approval and consent to participate

Not applicable.

## Consent for publication

Not applicable.

## Supplementary material

**Supplementary Figure 1.**
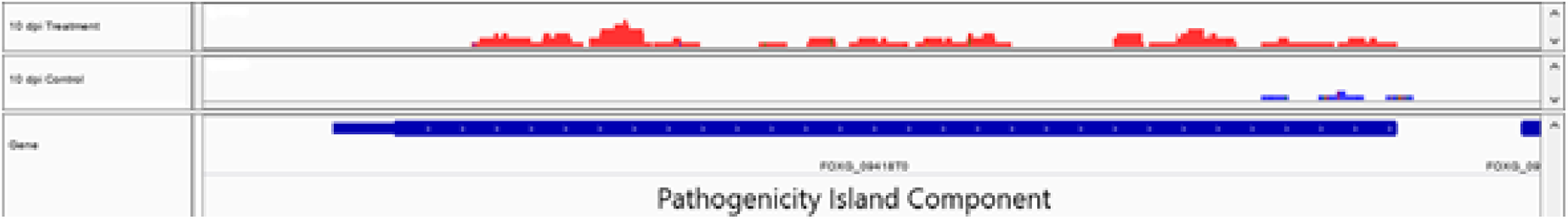
Track files visualization of the treatment strain JAGH3 (red tracks) at 10 dpi, its control (blue tracks) and the gene structure below in the genomic coordinates corresponding to a Pathogenicity Island Component in IGV Genome Browser.

## Supplementary Files

**2_dpi_treatment_genes_list.txt**

**10_dpi_treatment_genes_list.txt**

**Metabolic_routes_of_early_infection_at_2_dpi.xlsx**

**All_the_metabolic_routes_of_the_infection_dead_10_days.xlsx**

